# Altered Steroidogenesis in Response to Impaired Mitochondrial Protein Homeostasis

**DOI:** 10.64898/2026.11.02.763709

**Authors:** Mikhaela B. Slavin, Laurie P. Lee-Glover, Tian Zhao, Mezbah Uddin, Alvin Joselin, Rachel Wong, David S. Park, Davide Martino, Gerald Pfeffer, Timothy E. Shutt

## Abstract

Mitochondrial proteostasis is critical for organelle function, but how it influences mitochondrial steroidogenesis is not understood. We investigated the impact of impaired mitochondrial proteostasis on mitochondrial protein import and how this impacts cortisol production. We found that mild mitochondrial proteostatic stress increased cortisol synthesis whereas severe stress suppressed cortisol, revealing a biphasic steroidogenic response. Mild stress induced increases in cholesterol import and cortisol production in a manner dependent on STARD1-mediated cholesterol transport, which is the first and rate-limiting step in steroidogenesis. Severe stress reduced cortisol synthesis independently of STARD1 activity, indicating this inhibition occurs downstream in steroidogenesis. These findings reveal mitochondrial protein import as a regulatory node linking mitochondrial proteostasis to steroidogenic output. Furthermore, we observed increased cortisol synthesis in rotenone, A53T alpha-synuclein and PINK1-KO models of PD, suggesting that this response is relevant to PD-associated mitochondrial dysfunction. Together, our findings reveal a biphasic steroidogenic response to impairments in mitochondrial quality control with implications in human disease. Our work recognizes a previously unexplored branch of the mitochondrial stress response that enables cellular and physiological adaptations that can be detrimental if constitutively activated.

## INTRODUCTION

Mitochondria have multiple quality control pathways dedicated to maintaining protein homeostasis (proteostasis) (Song, Herrmann and Becker, 2021), which are required for these critical organelles to perform multiple functions beyond ATP production, including mediating apoptosis, calcium signaling, lipid metabolism, regulating innate immunity, and steroidogenesis. Among these pathways, mitochondrial protein import is required for the uptake of nuclear-encoded proteins that comprise ∼99% of the mitochondrial proteome. While impaired mitochondrial proteostasis triggers adaptive stress responses (Jin and Youle, 2013; Boos *et al*., 2019), we lack a complete understanding of these triggers and responses and their relevance to human disease.

Although mitochondrial disease has historically been associated with defects in oxidative phosphorylation (OXPHOS), novel pathogenic variants in certain mitochondrial proteins can cause disease without overt OXPHOS defects. For example, mitochondrial skeletal disorders due to impairments in proteins that mediate mitochondria proteostasis (*e.g.*, LONP1, HSPA9, IARS2, PAM16, AIFM1), are characterized by skeletal dysplasia and cataracts, and exhibit remarkable phenotypic heterogeneity that cannot solely be explained by impaired energy production (Zhao *et al*., 2020). These clinical presentations suggest a common mechanistic underpinning that is not energetic in nature, yet is clearly influenced by defects in protein homeostasis. In this context, elevated cortisol levels can potentially explain the skeletal abnormalities (Tack *et al*., 2016) and cataracts (Urban and Cotlier, 1986) seen in mitochondrial skeletal disorders, suggesting that altered mitochondrial steroidogenesis may be an underlying disease mechanism. Beyond rare mitochondrial skeletal disorders, mitochondrial proteostatic stress is also thought to contribute to mitochondrial dysfunction and neuronal cell death in Parkinson Disease (PD) (Di Maio *et al*., 2016; Franco-Iborra *et al*., 2018), where PD patients have documented increases in cortisol (Skogar *et al*., 2011; Costa *et al*., 2019).

Mitochondria play a critical role in adrenal steroidogenesis and the production of cortisol, a glucocorticoid that acts as the human body’s primary stress hormone. Importantly, the rate-limiting step of cortisol production is the import of cholesterol across the outer mitochondrial membrane (OMM), which is primarily mediated by Steroidogenic Acute Regulatory Protein (StAR/STARD1). Notably, STARD1 is negatively regulated by its own protein import efficiency into mitochondria, such that unimported STARD1 on the OMM is required for cholesterol import (Bose, Lingappa and Miller, 2002), while STARD1 import into the matrix renders it inactive for cholesterol import (Fig. 1A). While previous work shows that impaired mitochondrial protein import increases cholesterol import into mitochondria via STARD1 (Bose, Lingappa and Miller, 2002), the downstream effects on cortisol production have not been investigated.

**Figure 1.**
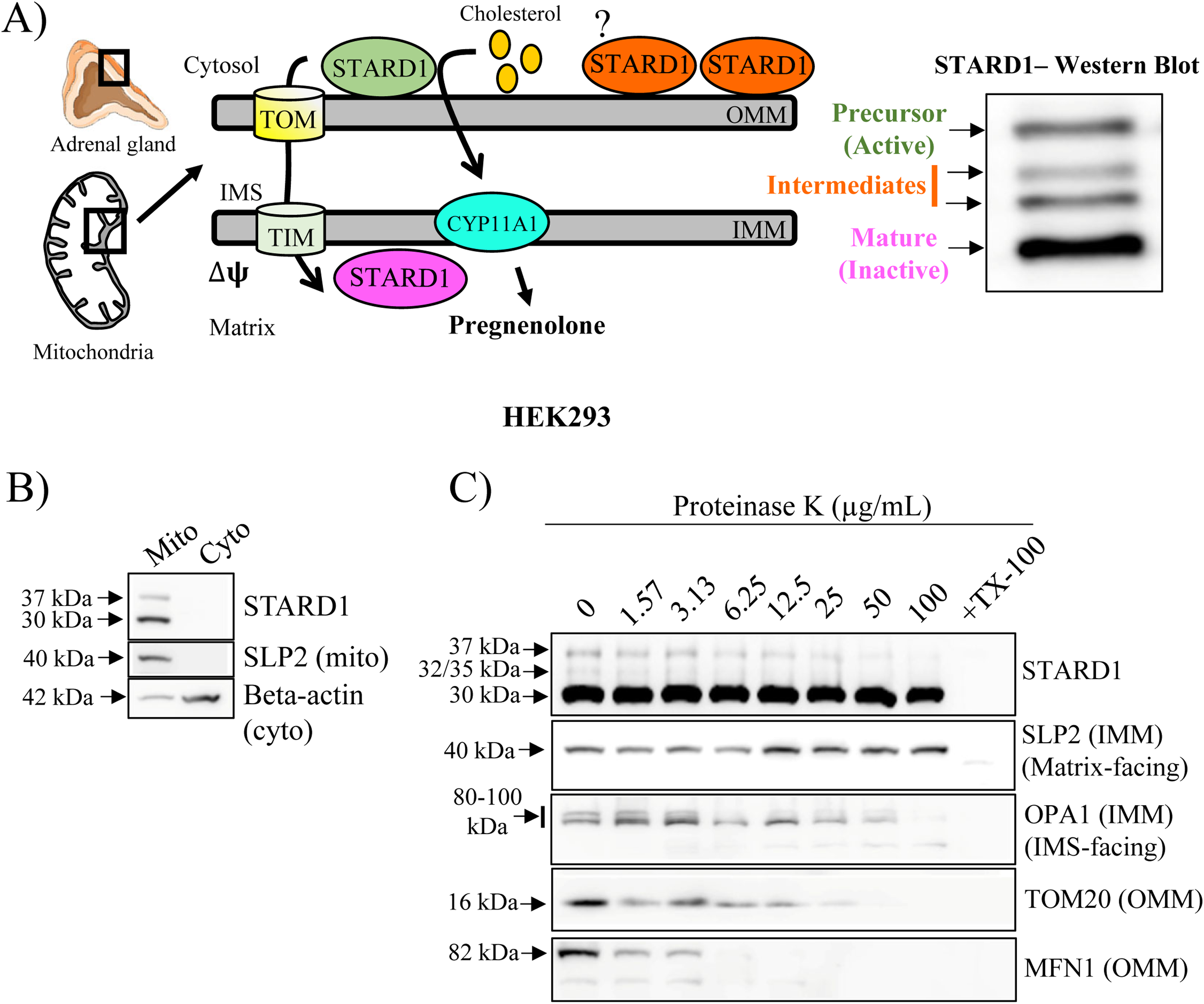
STARD1 processing regulates cholesterol import into mitochondria for steroidogenesis. A: Schematic of STARD1 activities at adrenal mitochondria to produce pregnenolone. The 37 kDa form is the full-length, active precursor form of STARD1, which is present on the OMM where it is exposed to the cytosol. The precursor transports cholesterol to the IMM enzyme CYP11A1, which synthesizes pregnenolone. When STARD1 is imported into the matrix, it is cleaved to its 30 kDa mature form and is steroidogenically inactive. Both forms can be visualized in a western blot, in addition to two intermediate forms that are detected. The function of the intermediate forms is unknown. ΔΨ, mitochondrial membrane potential. **B:** When transfected into HEK293 cells, STARD1 is detected in mitochondrial fractions. **C:** Protease protection assay in isolated mitochondria. Mitochondria were incubated with indicated concentrations of Proteinase K on ice for 15 minutes. Incubation with TX-100 was used as a control to permeabilize both membranes. Samples were then run on a western blot to probe for the indicated proteins. Cyto, cytosolic fraction; Mito, mitochondrial fraction.

Here, we uncover a novel biphasic response to cortisol production where mild perturbation of mitochondrial proteostasis impair mitochondrial import of STARD1 and stimulates cortisol production, whereas more severe mitochondrial proteostatic stress suppresses cortisol production. We also extend this novel stress response pathway to multiple PD models. Collectively, our findings establish mitochondrial steroidogenesis as a previously unappreciated component of the cellular response to mitochondrial stress and provide new insight into how mitochondria coordinate global physiological adaptation.

## RESULTS

### Visualizing STARD1 processing and import

To investigate the relationship between mitochondrial protein import and steroidogenesis, we set out to examine STARD1 processing as an indicator of its import efficiency into mitochondria.

However, the unprocessed form of STARD1 is not readily detectable in steroidogenic cells, which is speculated to be due to its short half-life (Epstein and Orme-Johnson, 1991; Granot *et al*., 2003). As reported by others, we also do not see unprocessed full-length STARD1 in steroidogenic H295R cells (Fig. S1A). Consistent with the rapid turnover of full-length STARD1, we were only able to detect a faint band only following combined treatment with dibutyryl-cAMP to increase STARD1 translation, FCCP to block mitochondrial import, and MG-132 to inhibit proteasomal degradation, followed by prolonged chemiluminescent exposure.

As an alternative approach to visualize STARD1 processing, transient expression of STARD1 is commonly used (Arakane *et al*., 1996; Bose, Lingappa and Miller, 2002). Thus, we transfected HEK293 cells with a STARD1 construct containing a C-terminal FLAG tag (Fig. S1B). In addition to the active, full-length (37 kDa) and the inactive, matrix-imported (30 kDa) forms, we also observed intermediate cleavage forms at 32 and 35 kDa (Fig. S1B) (Epstein and Orme-Johnson, 1991). To confirm the mitochondrial localization of these intermediate isoforms, we performed a protease protection assay on isolated mitochondria (Fig. 1B). Notably, the 35 kDa and 32 kDa bands were lost along with OMM markers (TOMM20 & MFN1), prior to loss of the inner mitochondrial membrane (IMM) (OPA1) and the matrix marker (SLP2). These results suggest that like the 37 kDa isoform, the 32 and 35 kDa STARD1 isoforms are primarily exposed to the cytosol, where they are likely capable of cholesterol transport.

### Impaired mitochondrial protein import reduces mitochondrial import of STARD1 and promotes mitochondrial cholesterol import

To confirm that STARD1 processing is sensitive to reduced mitochondrial protein import, we treated STARD1-transfected HEK293 cells with increasing doses of DNP (0-1000 µM for 24 hours), a protonophore that dissipates the mitochondrial membrane potential required for protein import into the mitochondrial matrix. DNP treatments reduced membrane potential in a dose-dependent manner, which we confirmed using TMRE staining and flow cytometry (Fig. 2A). Treatment with DNP yielded increases in the 37 kDa and intermediate forms with concomitant reductions in the 30 kDa forms (Fig. 2B). This work confirms that mitochondrial protein import regulates STARD1 processing.

**Figure 2.**
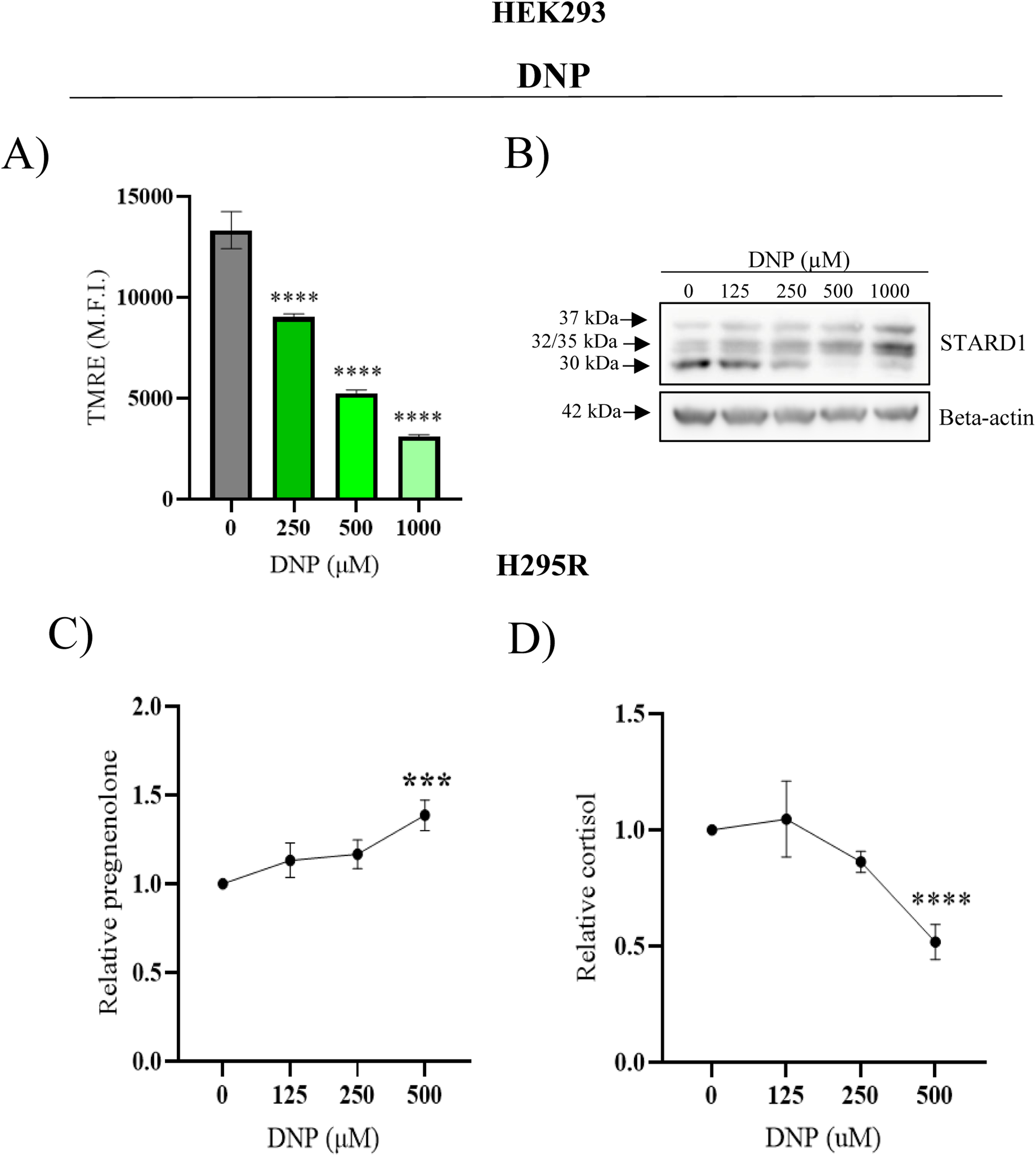
Effects of impaired mitochondrial protein import on STARD1 processing and steroidogenesis. A: HEK293 cells were treated with indicated concentrations of DNP and stained simultaneously with TMRE for 30 minutes. The Mean Fluorescence Intensity (M.F.I.) was measured with flow cytometry. Error bars represent the standard deviation of three biological replicates. A one-way ANOVA was used followed by Dunnett’s post hoc test to compare each dosage to the control group. **B:** STARD1 western blot from cells transfected with STARD1-FLAG and treated with DNP dosages for 24 hours. **C:** H295R cells were treated with indicated doses of DNP in combination with dibutyryl-cAMP and trilostane/abiraterone for 4 hours and relative pregnenolone levels from cell culture media were measured. **D:** Relative cortisol levels from cells treated as in A. For all panels, error bars represent the standard deviation from 3-4 biological replicates. ***p<0.001, ****p<0.0001, one-way ANOVA and Dunnett’s post hoc test comparing each dosage to the control group (0 µM).

Having confirmed that STARD1 processing is responsive to impaired mitochondrial protein import, we next investigated how DNP treatment impacts steroidogenesis. As HEK cells are not steroidogenic, we used the human adrenocortical cell line H295R and its monoclonal version, HAC15. These longstanding steroidogenic cell lines are considered the gold standard for studying cortisol steroidogenesis (Parmar, Key and Rainey, 2008; Odermatt, Strajhar and Engeli, 2016). We first set out to measure mitochondrial cholesterol import, which is the rate limiting step in steroidogenesis.

However, given that cholesterol is converted to pregnenolone nearly instantaneously upon its import into steroidogenic mitochondria (Kennedy, Charman and Karten, 2017), we measured pregnenolone production as a proxy for cholesterol import. To ensure accurate measurement of cholesterol import, the downstream conversion of pregnenolone to other steroid intermediates, including cortisol, was blocked using the inhibitors trilostane and abiraterone (Fig. S2A-C). Additionally, steroidogenesis was stimulated by treating cells with dibutyryl-cAMP (Fig. S2B,C). Consistent with the effects of DNP on STARD1 processing, we observed a dose-dependent increase in pregnenolone production, with increases of 18%, 20% and 34% at 125 µM, 250 µM and 500 µM of DNP, respectively (Fig. 2C) indicative of increased cholesterol import into mitochondria. Next, we measured cortisol production with DNP treatment. However, despite an increase in cholesterol import, which is typically the rate limiting step in steroidogenesis, we saw no stimulation of cortisol production. In fact, we found that the highest DNP dose (500 µM) actually reduced cortisol production by 48% compared to vehicle control (Fig. 2D), likely due to the fact that downstream steps in steroidogenesis depend on mitochondrial membrane potential (Prasad *et al*., 2012).

### Mitochondrial proteostatic stress reduces mitochondrial import of STARD1

Next, we set out to determine whether perturbing mitochondrial proteostasis is sufficient to impair STARD1 import. We first treated cells with CDDO, an inhibitor of the mitochondrial protease LONP1 (Lee *et al*., 2022). To confirm that this treatment induces proteostatic stress, we assessed the accumulation of protein aggregates using the Enzo Proteostat Aggresome Detection Kit. To this end, we found that a 4-hour treatment with 20 µM of CDDO resulted in a detectable increase in the fluorescence intensity of the proteostat dye, as quantified by flow cytometry (Fig. 3A). Next, we examined how CDDO impacts STARD1 import into mitochondria via immunoblotting. Consistent with reduced import of STARD1, we observed an increase in the levels of the full-length 37 kDa bands and the partially processed 32-35 kDa bands (Fig. 3B).

**Figure 3.**
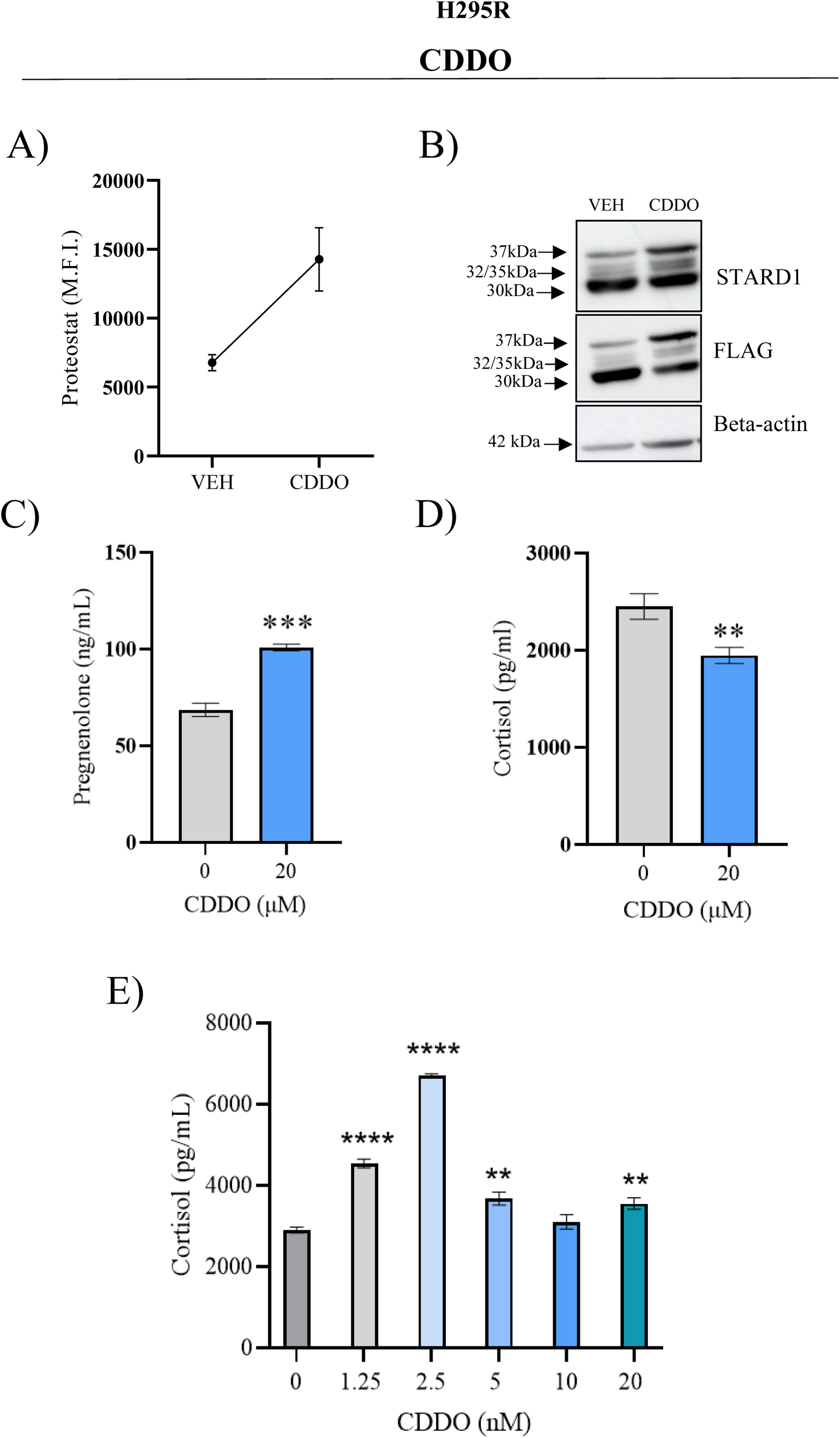
Impairments in mitochondrial proteostasis alter steroidogenesis. A: HEK293 cells were treated with 20 µM CDDO for 4 hours, then fixed and stained with the Proteostat dye. The Median Fluorescence Intensity (M.F.I.) was measured with flow cytometry. The graph is a representative experiment from two biological replicates. The error bars represent the standard deviation from three technical replicates. **B:** STARD1 and FLAG blots from transfected cells treated with 20 µM CDDO for 4 hours. Beta-actin was used as a blotting loading control. Blots are representative of three biological replicates. **C:** Pregnenolone and **D:** cortisol values from the media of H295R cells treated with 20 µM CDDO for 4 hours. For C and D: **p<0.01, ***p<0.001, unpaired t-test. **E:** Cortisol values from H295R cells treated with titrated doses of CDDO. For E: **p<0.01, ****p<0.0001, one-way ANOVA and Dunnett’s post hoc test comparing each dosage to the control group (0 µM). For C, D, and E, the graphs are a representative experiment of two biological replicates. Errors bars represent the standard deviation from three technical replicates.

### Mitochondrial proteostatic stress induces a biphasic response on cortisol production

To test the effects of proteostatic stress on steroidogenesis, H295R cells were then treated with CDDO. When we investigated cholesterol import, we noted that a 20 µM treatment with CDDO for 4 hours increased pregnenolone production by 47% compared to vehicle-treated cells (Fig. 3C).

However, despite increased cholesterol import, this high dose of CDDO treatment decreased cortisol production by 20% (Fig. 3D). Given this disparity, we performed a dose-response experiment with lower CDDO concentrations to induce relatively milder dysfunction. Intriguingly, we found that low doses of CDDO (1.25-20 nM) significantly increased cortisol production, with the 2.5 nM dose inducing a 2.3-fold increase (Fig. 3E).

### STARD1 is required for increased cholesterol import in response to impaired mitochondrial protein import and proteostasis

We next wanted to confirm whether the increased cholesterol import and cortisol production in response to proteostatic stress were dependent on STARD1. To this end, cells were treated with 22-OH-Chol, a membrane-permeable cholesterol analog that bypasses the need for STARD1-mediated import (Fig. 4). As expected, addition of 22-OH-Chol increased the rate of pregnenolone synthesis by 3.9-fold (Fig. 4A), as the enzymatic generation of pregnenolone by CYP11A1 is now rate-limiting, rather than STARD1 cholesterol import. However, addition of 500 µM DNP, a dose that stimulates pregnenolone production in control cells, did not further increase pregnenolone production in 22-OH-Chol treated cells (Fig. 4A). This finding indicates that the DNP increase of pregnenolone depends on cholesterol import.

**Figure 4.**
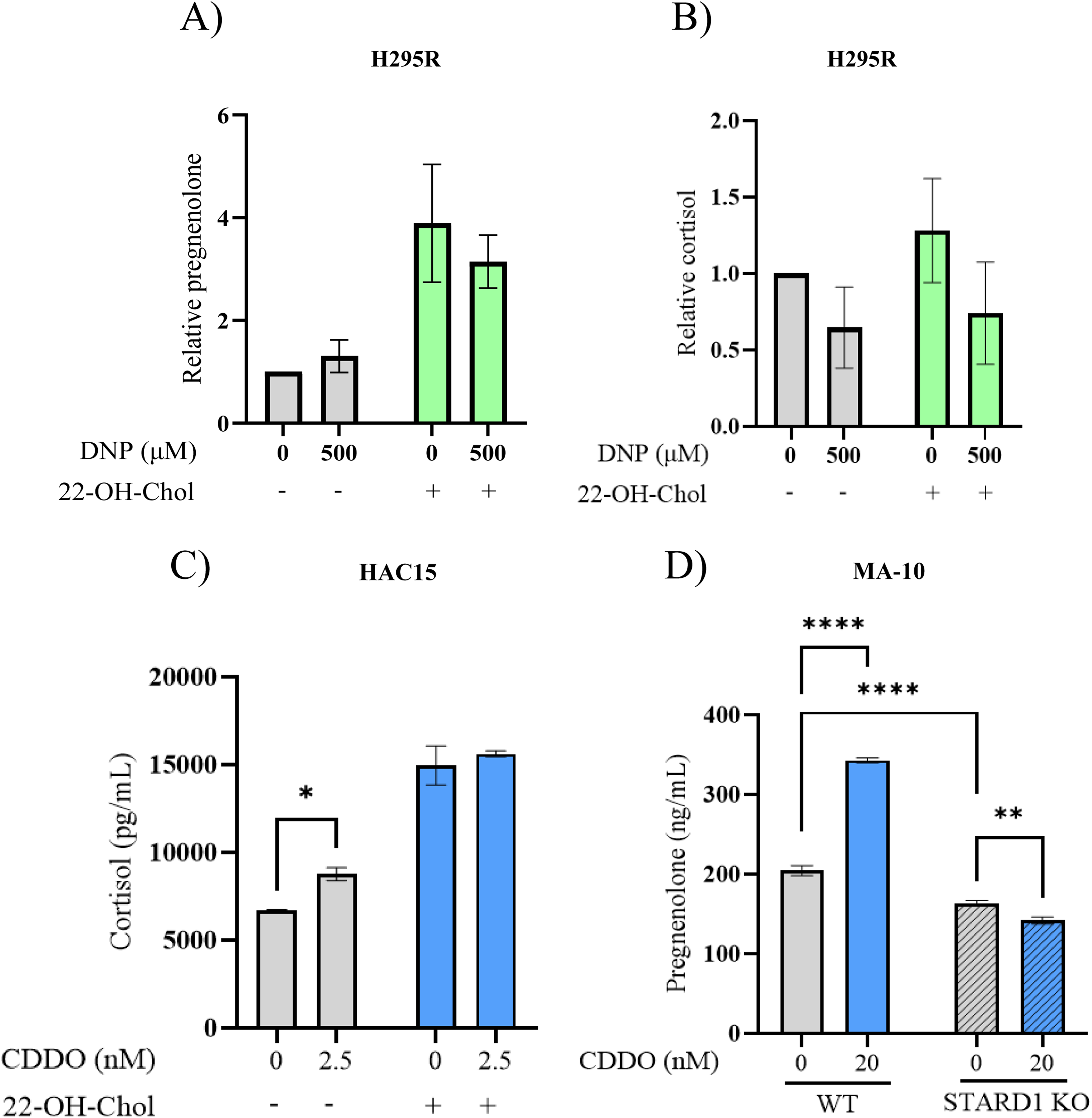
Increased cholesterol import is dependent on STARD1. Relative levels of pregnenolone (**A)** and cortisol **(B)** from media of H295R cells treated with either vehicle (DMSO) or 500 µM DNP with or without 5 µM 22-OH-Chol for 4 hours. Error bars represent the standard deviation from two (A) or four (B) biological replicates. No significance was found with a two-way ANOVA and Dunnett’s post hoc test. **C:** Cortisol measurements from HAC15 cells treated with 2.5 nM CDDO with or without 22-OH-Chol for 4 hours. **D:** Pregnenolone values from MA-10 WT and STARD1 KO cells treated with 20 nM CDDO. For C and D, error bars represent the standard deviation from three technical replicates, and it is a representative experiment from two biological replicates. *p<0.05, **p<0.01, ****p<0.0001, two-way ANOVA followed by Tukey’s post hoc test.

We then assessed cortisol production in cells treated with 22-OH-Chol. As expected, 22-OH-Chol also increased cortisol production (Fig. 4B). However, we observed that addition of 500 µM DNP, which reduces cortisol production in control cells, also decreased cortisol production in 22-OH-Chol treated cells, consistent with this dose of DNP impairing steroidogenesis downstream of STARD1 (Fig. 4B). Meanwhile, the low 2.5 nM dose CDDO that stimulates cortisol production no longer increases cortisol production in the presence of 22-OH-Chol (Fig. 4C). These findings indicate that low CDDO doses increase cortisol production by stimulating cholesterol import, while high CDDO inhibits an intermediate step in cortisol synthesis.

To confirm the role of STARD1 in mediating increased cholesterol import in response to proteostatic stress, we used WT and STARD1 KO MA-10 murine tumour Leydig cells that were generously donated by the Papadopoulos lab (Galano and Papadopoulos, 2022). First, we confirmed that STARD1 KO cells had the capacity to produce pregnenolone at levels comparable to WT cells when supplemented with 22-OH-Chol (Fig. S3). Next, we measured pregnenolone production in WT and KO MA-10 cells in response to 20 nM CDDO treatment. While CDDO treatment increased pregnenolone import by 68% in WT cells, STARD1 KO cells showed no increase in pregnenolone production (Fig. 4D), supporting the role of STARD1 in mitochondrial stress-induced cholesterol import.

### PD toxin rotenone impairs STARD1 import and alters steroidogenesis

Given the links between mitochondrial dysfunction and elevated cortisol in PD (Sherer *et al*., 2002; Skogar *et al*., 2011; Costa *et al*., 2019; Risiglione *et al*., 2020; Koçancı *et al*., 2026), we wanted to test whether the PD-relevant toxin rotenone, impacts STARD1 (Fig. 5). Rotenone is a potent inhibitor of mitochondrial Complex I (Ravipati *et al*., 2026) that leads to neurodegeneration in nigral dopaminergic neurons and is used to model experimental parkinsonism (Martinez and Greenamyre, 2012). Treating cells with rotenone for 24 hours caused dose-dependent increases in full-length and intermediate forms of STARD1 (Fig. 5A). We then examined how these doses impact mitochondrial membrane potential and found that rotenone treatments reduced membrane potential by approximately 40% (Fig. 5B), Thus, STARD1 import is reduced by rotenone, which also impacts membrane potential.

**Figure 5.**
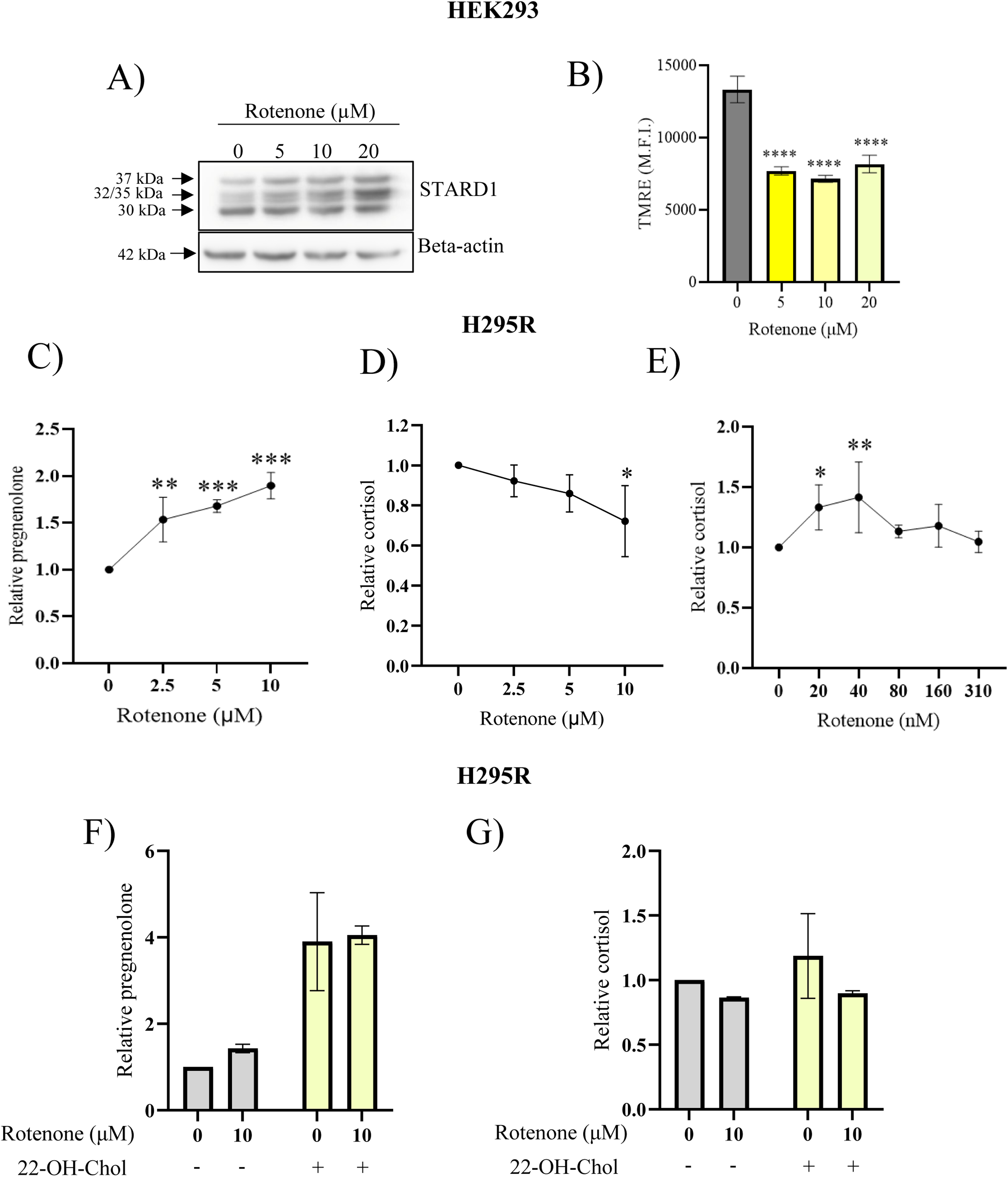
Effect of PD-relevant toxin Rotenone on STARD1 import and steroidogenesis. A: HEK293 cells were treated with indicated concentrations of Rotenone and stained simultaneously with TMRE for 30 minutes. The Mean Fluorescence Intensity (M.F.I.) was measured with flow cytometry. Error bars represent the standard deviation of three biological replicates. A one-way ANOVA was used followed by Dunnett’s post hoc test to compare each dosage to the control group. **B:** Cells were transfected with STARD1 and treated 24 hours later with indicated concentrations of Rotenone for 24 hours. STARD1 processing was assessed by western blotting, with beta-actin as a loading control. Blots are representative of three biological replicates. Relative **C:** pregnenolone and **D,E:** cortisol levels from H295R cells treated for 4 hours with indicated doses of Rotenone. *p<0.05, **p<0.01, ***p<0.001, ****p<0.0001, one-way ANOVA and Dunnett’s post hoc test comparing each dosage to the control group (0 µM). Error bars represent the standard deviation of three biological replicates. Relative **G:** pregnenolone and **H:** cortisol levels from H295R cells treated with 10 µM Rotenone with or without 5 µM 22-OH-Chol for 4 hours. Errors bars represent the standard deviation from two biological replicates.

We next tested if rotenone impacts mitochondrial cholesterol import and cortisol production in H295R cells (Fig. 5C-E). Notably, rotenone treatment induced dose-dependent increases in pregnenolone production by 53%, 68%, and 90% at the 2.5 µM, 5 µM, and 10 µM doses, respectively (Fig. 5C). When we then investigated cortisol, we found that the initial experiments using concentrations of rotenone that stimulated pregnenolone production led to reduced cortisol synthesis (Fig. 5D). However, when we tested a lower range of rotenone concentrations, we found that 20-40 nM rotenone increased cortisol levels by 30-40% (Fig. 5E). Notably, pregnenolone was also increased with 40 nM rotenone treatment (Fig. S4).

Next, we took advantage of 22-OH-Chol to understand whether the rotenone-induced increases in pregnenolone and cortisol were dependent on cholesterol import (Fig. 5F,G). As with DNP treatment, a rotenone dose that induces cholesterol import did not further increase pregnenolone production in the presence of 22-OH-Chol (Fig. 5F). Meanwhile, the same dose of rotenone prevented the 22-OH-Chol stimulation of cortisol production (Fig. 7G). These findings show that the stimulatory effects of rotenone on pregnenolone production depend on cholesterol import, while the inhibitory effects of rotenone on cortisol production are downstream of STARD1.

**Figure 6.**
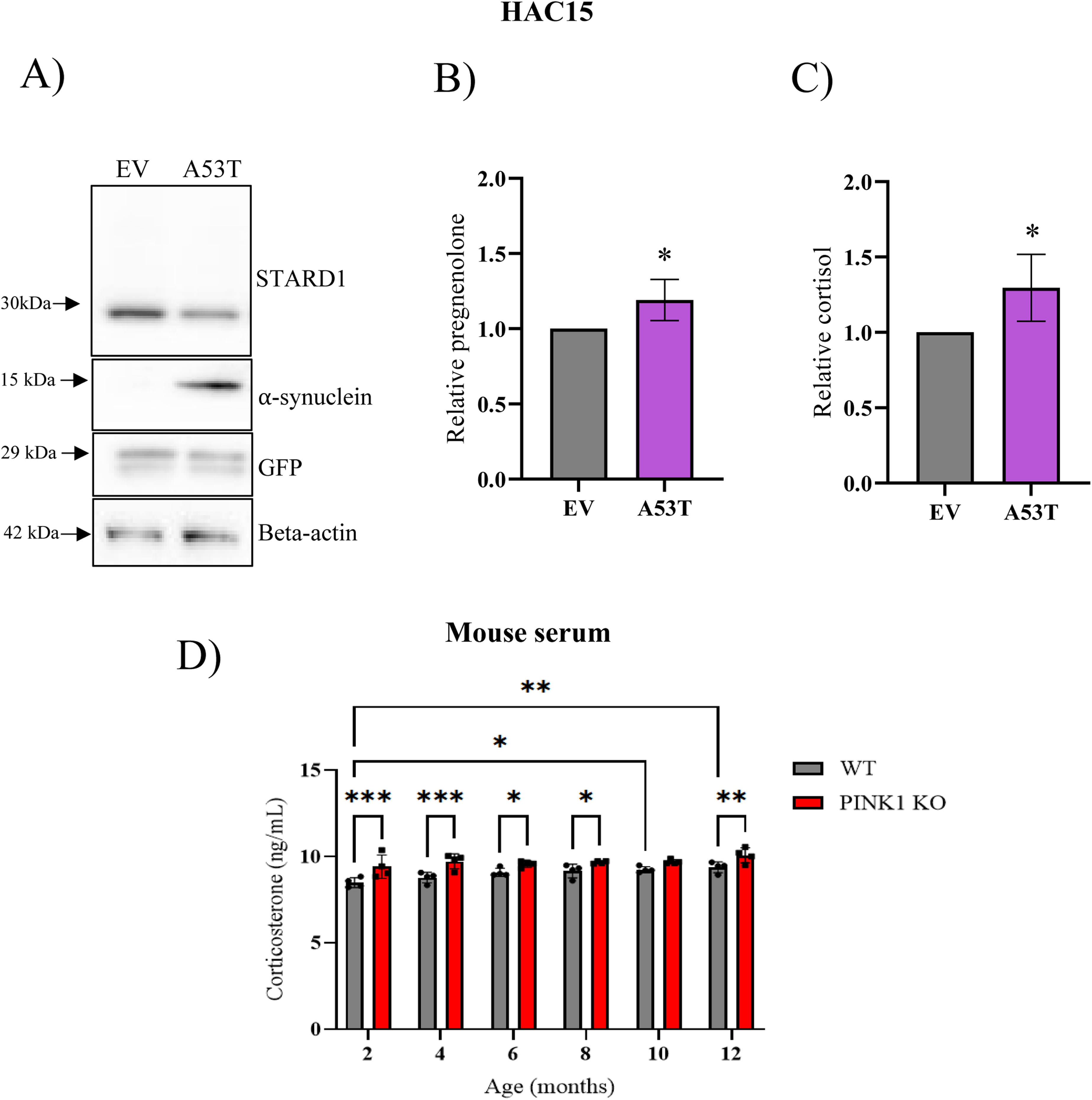
Upregulated steroidogenesis from biphasic model extends to PD. A: Western blots from HAC15 cells over-expressing alpha-synuclein A53T or empty vector (EV). Blots are endogenous STARD1, alpha-synuclein, and GFP, and are representative of four biological replicates. Beta-actin was used as a loading control and GFP was used to confirm expression of both vectors. Relative **B:** pregnenolone and **C:** cortisol levels from EV and A53T cells. Error bars represent the standard deviation from four biological replicates. For B and C: *p<0.05, unpaired t-test to compare both groups. **D:** Corticosterone values from mouse serum of WT and PINK1 KO mice at the ages indicated. Errors bars represent the standard deviation across animals (n=4). *p<0.05, **p<0.01, ***p<0.001, two-way ANOVA followed by Tukey’s post hoc test.

**Figure 7.**
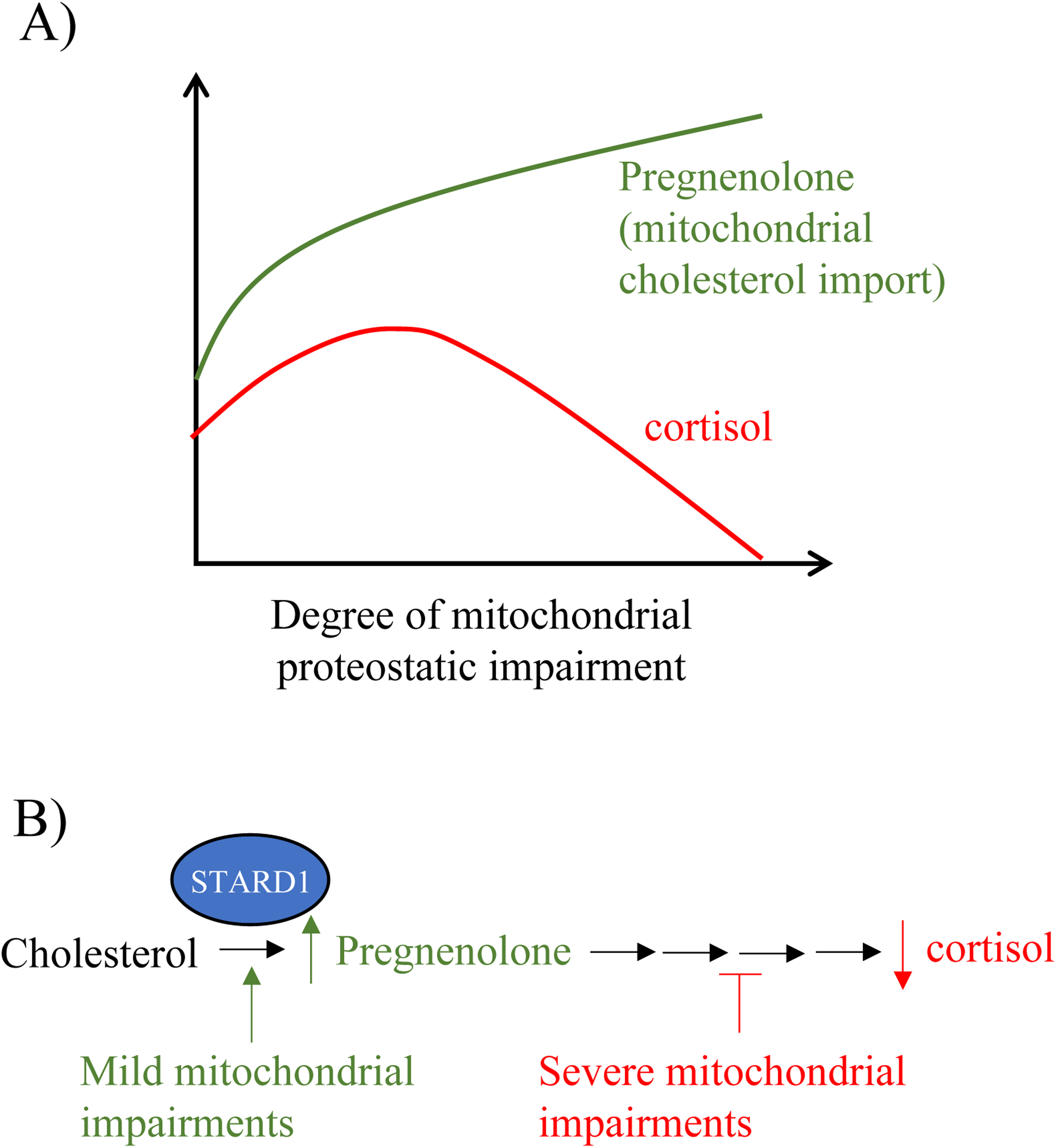
Steroidogenesis is regulated in a biphasic manner in response to mitochondrial stress. A: Diagram illustrating our biphasic steroidogenesis model. At lower levels of mitochondrial stress, both cholesterol import (pregnenolone) and cortisol synthesis are stimulated. At higher levels of dysfunction, there is an uncoupling of these processes as cholesterol import is still increased while cortisol production is reduced. **B:** Diagram depicting where in the pathway mitochondrial dysfunction may be influencing steroidogenesis. Upon mild mitochondrial impairments, the increase in pregnenolone is likely driven by an increase in STARD1-mediated cholesterol import, driven by the altered mitochondrial import and processing of STARD1. The reductions in cortisol upon high levels of mitochondrial stress are independent of STARD1-mediated cholesterol import, likely inhibiting steroidogenic steps downstream of cholesterol import.

### Expression of PD-related A53T -synuclein enhances steroidogenesis

To investigate whether genetic causes of PD are also relevant to STARD1 and steroidogenesis, we took advantage of the A53T mutation in the alpha-synuclein protein that causes PD (Choubey *et al*., 2011), and which is known to contribute to mitochondrial dysfunction, in part, by impairing mitochondrial protein import (Di Maio *et al*., 2016; Vasquez *et al*., 2024). As H295R cells are a mixed population (Wang and Rainey, 2012), which could introduce unwanted variability upon clonal expansion, we used HAC15 cells, which are a monoclonal derivative of H295R (Wang and Rainey, 2012). To this end, HAC15 cells were stably transduced with a lentivirus construct containing a GFP open reading to express A53T alpha-synuclein, or a GFP only empty vector (EV) virus as a control (Fig. 6). Notably, cells expressing A53T exhibited a reduction in the 30 kDa form of STARD1 (Fig. 6A), consistent with reduced STARD1 import. When we interrogated steroidogenesis in these cells, we noted that even in the absence of dibutyryl-cAMP stimulation, A53T expression induced a 19% increase in pregnenolone (Fig. 6B), alongside a 30% increase in cortisol synthesis (Fig. 6C).

### Increased stress hormone levels in PINK1 KO mice

To investigate if steroidogenesis is impacted in an animal model of PD, we used PINK1-KO mice, as mutations in PINK1 cause PD (Gandhi *et al*., 2012). We measured levels of corticosterone, the murine equivalent of cortisol, in blood samples from WT and KO mice obtained across six ages (Fig. 6D). In accordance with our cellular models, we observed a significant increase in corticosterone from PINK1 KO samples at 2, 4, 6, 8 and 12 months of age with a trend of increased corticosterone at 10 months. We also observed an effect of age in WT mice, with increased corticosterone at 10 and 12 months of age relative to the 2 month-old animals.

## DISCUSSION

We define a novel mechanism whereby cortisol synthesis is stimulated upon perturbations in mitochondrial protein import of STARD1 that can be induced through mild impairment of mitochondrial proteostasis. However, upon more severe impairment, cortisol synthesis is inhibited downstream of cholesterol import. This biphasic response of cortisol production (Fig. 7A) represents a previously uncharacterized branch of the mitochondrial stress response and has important implications for human diseases such as PD. Specifically, we demonstrate that pharmacologic and genetic models of PD recapitulate elevated cortisol levels. These findings are consistent with a model where impaired mitochondrial protein import senses mitochondrial dysfunction and elicits numerous stress responses, which we show here also includes increased cortisol production.

### Mitochondrial stress elicits a biphasic cortisol response

Historically, mitochondrial dysfunction in human disease has been primarily attributed to severe defects in OXPHOS. Similarly, alterations in steroidogenesis associated with mitochondrial dysfunction have largely been studied in the context of adrenal insufficiency (Corkery-Hayward and Metherell, 2023). Consistent with earlier studies demonstrating impaired steroidogenesis under conditions of substantial mitochondrial dysfunction (King and Stocco, 1996; King *et al*., 1999; Artemenko *et al*., 2001), we also observed reduced cortisol synthesis in response to severe mitochondrial stress. However, our results reveal an unrecognized response to milder mitochondrial dysfunction: the stimulation of glucocorticoid steroidogenesis.

A similar biphasic steroidogenic response, also referred to as hormesis (Mattson, 2008), has been described previously with a wide range of chemicals, including rotenone, which stimulate steroidogenesis under lower concentrations and inhibit it under higher concentrations (Zhao *et al*., 2012; Štefunková *et al*., 2023; Mathis *et al*., 2024). However, the mechanism beneath this biphasic response was unknown. In our findings, we expand this biphasic response to proteostatic stress, and provide a mechanistic basis for this phenomenon (Fig. 7B). Specifically, we show that increased cortisol production at lower doses/stresses is caused by impaired mitochondrial import of STARD1 and increased cholesterol import, which is the rate limiting step of steroidogenesis. Meanwhile, inhibition of cortisol production at higher doses/stresses is caused downstream of STARD1 import and its role in cholesterol transport. One potential mechanism for this downstream inhibition is the impaired activity of 3ß-hydroxysteroid dehydrogenase 2 (3ß-HSD2), the enzyme that catalyzes the conversion of pregnenolone to 17-hydroxyprogesterone, a steroid intermediate in cortisol synthesis. Notably, 3ß-HSD2 activity requires the mitochondrial proton gradient, such that mitochondrial depolarization reduces 3ß-HSD2 activity (Prasad *et al*., 2012). Additionally, increased ROS, which can occur upon inhibition of the electron transport chain (Chen *et al*., 2007), may also contribute to reduced cortisol production. Consistent with this possibility, disturbances in adrenal redox homeostasis are implicated in adrenocortical insufficiency (Meimaridou *et al*., 2012; Prasad *et al*., 2014). Delineating the mechanisms underlying the inhibition of steroidogenesis under conditions of severe mitochondrial dysfunction will be relevant to a variety of mitochondrial diseases where reductions in cortisol synthesis are observed.

### Cortisol upregulation in disease

Our novel findings that mild mitochondrial stress can stimulate cortisol production, and the elucidation of the underlying mechanism, have implications for diseases. Given the multiple functions performed by mitochondria and the heterogeneity of patient phenotypes associated with mitochondrial disease, it is tempting to speculate that impairment of different mitochondrial functions may lead to different pathologies. Similarly, different types of mitochondrial dysfunction likely elicit different stress responses that contribute to disease heterogeneity. In this context, increased cortisol synthesis may be relevant in mitochondrial disorders characterized by more mild impairments in mitochondrial quality control. Although acute elevations in cortisol can have transient benefits to mitochondrial function (Du *et al*., 2009), constitutive elevation of cortisol can be detrimental, as it can cause cataracts (Urban and Cotlier, 1986), impair neuronal development (Fukumoto *et al*., 2009), and inhibit skeletal growth (Tack *et al*., 2016), which are phenotypes exhibited in mitochondrial skeletal disorders (Zhao *et al*., 2020).

Beyond rare mitochondrial diseases, mitochondrial dysfunction also contributes to other diseases such as PD. Notably, elevated levels circulating cortisol are documented in PD patients (Hartmann *et al*., 1997; Djamshidian *et al*., 2011; Skogar *et al*., 2011; Costa *et al*., 2019), and are associated with poorer functional outcomes (Soares *et al*., 2019). Consistent with these clinical observations, we found increases in cortisol synthesis in multiple PD-relevant models. Notably, the PD-inducing chemical rotenone reduced membrane potential, impaired STARD1 import and increased cholesterol import. Meanwhile, the genetic perturbations, A53T and PINK1-KO, are also both linked to mitochondrial protein import. A53T alpha-synuclein is documented to impair mitochondrial protein import (Vasquez *et al*., 2024), while mitochondrial protein import efficiency is impaired in cell models harbouring a loss of PINK1 and in PD patient-derived cells with PINK1 mutations (Jacoupy *et al*., 2019). Together, these findings suggest that enhanced glucocorticoid production may be a shared feature across models of PD-associated mitochondrial dysfunction, driven by impairments in mitochondrial protein import to elicit an adaptive steroidogenic response.

### Mitochondrial protein import is a sensitive signaling node to stimulate cortisol synthesis

While mitochondrial protein import is recognized to act as a sensor of mitochondrial function to influence both organelle and cellular homeostasis (Lee-Glover and Shutt, 2024), our work here linking elevated cortisol to impaired mitochondrial protein import identifies a novel arm of this response. One of the earlier examples of mitochondrial protein import sensing mitochondrial dysfunction is the mitochondrial Unfolded Protein Response (UPR^mt^), a protective transcriptional mechanism that has been well-described in *C. elegans* (Naresh and Haynes, 2019). While transient activation of the UPR^mt^ in *C. elegans* is beneficial and has even been shown to promote organismal longevity (Lionaki *et al*., 2022), prolonged and/or excessive UPR^mt^ activation is associated with deleterious effects (Lin *et al*., 2016). Meanwhile, in mammalian models, the DELE1/HRI arm of the integrated stress response (ISR) also listens to mitochondrial import (Bi *et al*., 2024). While both the UPR^mt^ and ISR detect mitochondrial dysfunction at the cellular level, they can also lead to systemic responses (Zhang *et al*., 2018; Forsström *et al*., 2019). The novel mitochondria-steroidogenesis axis thus joins these other stress-response pathways as a new mechanism that is stimulated upon mitochondrial protein import stress to drive cellular and physiological adaptation.

### Limitations

One limitation of our study is the inability to directly visualize active full-length STARD1 in steroidogenic cells, an issue that has been noted in other studies as well (Granot *et al*., 2007). Instead, we rely here on making associations between STARD1 processing in non-steroidogenic cells with the dynamics of cholesterol import and cortisol synthesis in steroid-producing cells. Nonetheless, we see consistent correlations across multiple treatments and models that support our model. Meanwhile, the link between PD, which is a neurodegenerative disease, and the adrenal steroidogenic lines we used, which are well-characterized and established models to study steroidogenesis, may not be immediately evident. However, it is worth noting that glucocorticoids can also be synthesized in non-adrenal tissues, including the brain, lung, skin, placenta, adipose tissue, and spleen (Slominski *et al*., 2020). The brain is particularly interesting given its capacity for neurosteroidogenesis, and the fact that cells in the CNS such as microglia, neurons and oligodendrocytes, express steroidogenic machinery including STARD1 (Sierra, 2004). Thus, the adaptive steroidogenic response characterized in our study likely extends beyond the adrenal glands to other cortisol-producing tissues. Future studies should investigate whether the biphasic steroidogenic response to mitochondrial dysfunction observed in our study also extends to the brain and neurosteroid production.

## Conclusion

Our findings identify altered mitochondrial quality control as a regulator of cortisol production, revealing a biphasic response to low and high levels of mitochondrial stress. We show that steroidogenesis is sensitive to disruptions in mitochondrial function beyond energetic impairment, particularly disruptions in mitochondrial protein import and proteostasis. Under mild stress, increased cortisol synthesis is caused by impairments in STARD1 import, highlighting mitochondrial protein import as a sensitive signaling node that senses impairments in proteostasis and engages an adaptive steroidogenic response. This steroidogenic axis represents a previously unrecognized branch of the mitochondrial stress response that likely contributes to cellular and physiological adaptation. This work has implications for understanding the pathological mechanisms in PD, where mitochondrial dysfunction has long been recognized. Overall, we highlight steroidogenesis as an important, yet underexplored consequence of disrupted mitochondrial quality control that contributes to disease pathogenesis.

## Author contributions

Designed and/or performed and analyzed experiments: MBS, LPLG, TZ, MU, AJ, TES. Drafted manuscript and figures: MBS, TES. Funding acquisition: DM, GP, TES. Supervision: DP, DM, GP, TES. All authors discussed, commented on and consented authorship on the manuscript.

## Acknowledgements

We would like to acknowledge and thank Dr. Vassilios Papadopoulos and Melanie Galano at the University of Southern California for kindly donating the MA-10 wild-type and STARD1 KO cells.

## MATERIALS AND METHODS

### Cell culture

HEK293 cells were cultured in DMEM with 10% heat-inactivated fetal bovine serum (Gibco, A5670701). H295R cells were cultured in DMEM:F12 (Gibco 11320033) supplemented with ITS+ (Corning, 354352) and 2.5% Nu serum (Corning, 355100). HAC15 cells (ATCC, CRL-3301) cells were cultured in DMEM/F12 with ITS+ (Corning, 354352) and 10% heat inactivated cosmic calf serum (Cytiva, SH30087.03HI) on plates coated with poly D-lysine (Gibco, A3890401). MA-10 cells were cultured in DMEM:F12 with 1% of sodium bicarbonate 7.5% solution (Gibco, 25080094), 1% sodium pyruvate (100 mM) (Gibco, 11360070), 2.5% heat inactivated horse serum, (Gibco, 16050122) and 5% heat-inactivated fetal bovine serum. All cells were maintained at 37°C and in 5% CO_2_.

Cells were treated with CDDO (Bardoxolone) (AdooQ Bioscience, A11324-10), rotenone (Enzo Life Sciences, ALX-350-360-G005), and DNP (2,4-dinitrophenol) (Millipore Sigma, 34334) at indicated concentrations. Other drug treatments included: dibutyryl-cAMP (Santa Cruz Biotechnology, sc-201567A) at 50 µM; trilostane (AdooQ Bioscience, A10949-50) and abiraterone (AdooQ Bioscience, A10020-50) at 1 µM each; 22-OH-Chol (22(R)-hydroxy cholesterol) (Cayman Chemicals, 89355) at 5 µM; and FCCP (Enzo Life Sciences, BML-CM120-0010) at 10 µM.

### Transient transfections

HEK293 cells were seeded at 4 x 10^5 cells per well in a 6-well dish. Cells were transfected 24 hours later with 2.5 µg STARD1 plasmid DNA using Lipofectamine 3000 (Invitrogen, L3000001). Cells were harvested 48 hours post-transfection.

### Western blotting

Cells were harvested and lysed in RIPA buffer (Thermo Scientific, 89901) with 1x protease inhibitor cocktail (BioVision, K272-1). The soluble protein fraction was collected as a supernatant after centrifugation at 14 000 x g for 30 minutes. Protein concentration was determined using bicinchoninic acid assay (Thermo Scientific, 23223 and 23224). Unless otherwise indicated, 50 µg of protein was loaded per well. Samples were mixed with 4 x Laemmli buffer (Bio-Rad, 1610747) with 10% beta-mercaptoethanol and boiled for 5 minutes at 95°C. Proteins were separated on SDS-containing 10 or 12% polyacrylamide gels along with Precision Plus Protein All Blue Prestained Standards (Bio-Rad, 1610373) as molecular weight markers. Protein was transferred to a PVDF membrane overnight.

Blocking and antibody incubations were performed in 5% milk in TBS-T. Antibodies used are listed in Table S1. Bands were visualized by chemiluminescence using Clarity Western ECL Substrate (Bio-Rad, 1705061) or Super Signal West Femto (Thermo Scientific, 34095).

### Measurement of mitochondrial membrane potential

Cells were seeded into 6-well plates and treated with either rotenone or DNP as indicated. To measure mitochondrial membrane potential, cells were incubated with 50 nM TMRE for 30 minutes. As a control, 10 µM FCCP was added to depolarize mitochondria. Cells were then washed with PBS 4 times before being harvested in complete media. Cells were analyzed on an Attune NxT Flow Cytometer (Invitrogen) using the yellow laser (561 nm) and fluorescence intensity was recorded using the YL1 (585/16 nm) filter.

### Measurement of cellular proteostasis

Cells were seeded into 6-well plates, and after 24 hours were treated with 20 µM CDDO for 4 hours. Once cells were collected, aliquots of 1 x 10^6 cells per condition were used for subsequent fixation and staining. Cells were washed with PBS then spun down and fixed in suspension with 4% paraformaldehyde solution in FACS buffer (3% fetal bovine serum, 2 mM EDTA in PBS) at room temperature for 30 minutes. Subsequently, cells were washed once with PBS and permeabilized with permeabilization buffer (0.5% TX-100 in FACS buffer) for 30 minutes on ice. After one PBS wash, cells were suspended in staining buffer, composed of the proprietary Proteostat Aggresome Detection Reagent diluted 1/7000 in FACS buffer, and incubated in the dark for 30 minutes (Proteostat Aggresome Detection Kit, Enzo, ENZ-51035). Cells were then immediately analyzed (50,000 cells/condition) using a Sony LE-SH800 Cell Sorter in Analysis mode. The 488 nm laser was used for excitation, and emission was acquired using the FL3 (600/60) filter.

### Mitochondrial protease protection assay

A crude mitochondrial fraction was isolated from cultured cells. Two 10 cm dishes of HEK293 cells transfected with STARD1 were harvested and resuspended in hypotonic buffer from Mitochondrial Isolation Kit for Cultured Cells (Thermo Scientific, 89874). Cells were mechanically lysed in a dounce homogenizer until only 50% of cells were broken open, as assessed with trypan blue under the microscope. The mixture was spun at 700 x g for 10 minutes to pellet intact cells and nuclei. The post-nuclear supernatant was removed and spun again at 9000 x g for 15 minutes. The supernatant (cytosol) was removed and the pellet, containing mitochondria, was resuspended in 30 µL mitochondrial isolation buffer (220 mM mannitol, 68 mM sucrose, 80 mM KCl, 0.5 mM EGTA, 2 mM MgAc_2_, and 10 mM HEPES, pH 7.4). A volume of 1 µL of the mitochondrial fraction was taken and diluted 1:60 for protein concentration measurement with BCA.

For the protease protection assay, 30 µg of mitochondria per reaction were incubated on ice for 15 minutes in the presence of increasing concentrations of Proteinase K (VWR, CA97062-238) (1.6-100 µg/mL). As a complete digest control, 1% TX-100 was included to solubilize both membranes.

PMSF was added to a final concentration of 4 mM to inhibit proteinase K. Samples were mixed with loading buffer and boiled before being run on an SDS-PAGE gel and visualized with western blotting. **Pregnenolone and cortisol measurements from cell culture media**

As steroid hormones are cell permeable, pregnenolone and cortisol were measured from cell culture media. Steroidogenic cells were seeded in 6-well or 24-well plates. Cells were rinsed with PBS and complete culture medium was replaced with serum-free medium supplemented with 50 uM dibutyryl-cAMP (Santa Cruz Biotechnology, sc-201567A). When media was collected for pregnenolone measurements, media also included 1 µM each of trilostane (AdooQ Bioscience,

A10949-50) and abiraterone (AdooQ Bioscience, A10020-50) to prevent conversion of pregnenolone to other downstream steroids. Collected media was spun at 1000 x g for 10 minutes, then the supernatant was removed and stored at -80°C. Cortisol ELISA kits (Cayman Chemicals, 500360) and pregnenolone ELISA kits (IBL America, IB59107) were used to quantify steroid concentrations following the manufacturer’s instructions. If required, samples were diluted between 1:4 to 1:50 in serum-free media. A sample of growth media was analyzed as a negative control, and if necessary, used to subtract any background signal. In cases where pregnenolone or cortisol levels were compared between different cell lines (EV vs. A53T transduced HAC15 cells), an additional normalization step using protein levels was taken to adjust for any discrepancies in seeding densities. Following media collection, cells in the well were harvested, lysed and protein concentration was quantified. Relative protein concentration was calculated relative to control cells, and this factor was used to normalize the measured steroid levels from each well.

### Lentiviral transduction

HEK293 cells were seeded at 2.5 x 10^6 cells per 10 cm dish. Cells were transfected 24 hours later with 6 µg transfer vector, 5 µg packaging vector psPAX2 (Addgene plasmid #12260) and 2.5 µg envelope vector pMD2.G (gift from Didier Trono) using calcium phosphate transfection. The following transfer vectors were used: pLV-empty vector (EV) and pLV-alpha-synuclein [A53T] which were designed and ordered through VectorBuilder. pLV-alpha-synuclein [A53T] (VB220719-1381wwd) encodes alpha-synuclein containing the mutation c.157G>A (p.Ala53Thr) expressed under the EF-1A promoter. For selection, an eGFP and a puromycin resistance gene were expressed under a separate promoter, split by a T2A sequence. The pLV-EV vector was used as a transduction control. Media was replaced with fresh media containing 1 mM sodium butyrate 16 hours later. Media containing virus was collected at 48 hours, 72 hours and 96 hours post-transfection. Virus supernatant was passed through a 0.45 µm sterile filter to remove any cells and if not used right away, virus was frozen at -80°C.

For lentiviral transduction, virus was added along with 10 µg/mL hexadimethrine bromide (Polybrene) (Sigma-Aldrich, 107689). Two days after transduction, antibiotic selection was performed using puromycin (Sigma-Aldrich, P8833) in the presence of 10 µM verapamil to inhibit the P-glycoprotein (Bello-Reuss *et al*., 2000; Yang *et al*., 2019). Antibiotic-containing media was refreshed every 2-3 days until selection was complete.

### Stress hormone measurements from mouse serum

WT and PINK1-KO C57BL/6N mice were used in collaboration with Dr. David S. Park’s lab at the University of Calgary. Blood samples were collected in the morning from a tail snip, then spun at 2000 x g for 5 minutes at room temperature to separate serum from the hematocrit. Serum samples were then stored at -80°C until analysis by ELISA. To measure corticosterone levels in samples, the Bio Vision Cortisol Mouse/Rat/Human ELISA kit (K7430-100) was used. Samples were assayed in the provided 96-well plate and measured at 450 nm according to the kit instructions. Raw sample values were plotted against a standard curve from which the raw corticosterone measurements in ng/mL were extrapolated.

### Statistical analysis

Data were analyzed using GraphPad Prism 8.0 software and values are reported as means ±SD. Comparisons between two groups were carried out with unpaired Student’s t-tests. A one-way ANOVA was used in the comparison of multiple drug dosages to the vehicle-treated group. A 2-way ANOVA was used in the comparisons of multiple variables. The ANOVA was followed by the appropriate post hoc test where necessary. Statistical significance was set at P < 0.05. Significance was stated as follows: *p<0.05, **p<0.01, ***p<0.001, ****p<0.0001.

---

**Figure S1.** Visualization of STARD1 in H295R and transfected HEK293 cells. A: Western blots of endogenous STARD1 in H295R cells treated with 50 µM dibutyryl-cAMP and 10 µM MG-132 for 6 hours, with the addition of 10 µM FCCP for the last 4 hours. Both STARD1 blots are from the same gel but with the bottom blot being subjected to a longer chemiluminescent exposure. **B:** Western blots of exogenous STARD1 and FLAG from HEK293 cells transiently transfected with STARD1-FLAG. Beta-actin was used as a loading control.

**Figure S2.** Detection of steroids from H295R cell culture media. A: Diagram illustrating the initial steps of steroidogenesis and the site of action of the inhibitors. Levels of **B**: pregnenolone and **C:** cortisol in media collected from H295R cells treated with 50 µM dibutyryl-cAMP and 1 µM each of the inhibitors trilostane and abiraterone for 4 hours. Error bars represent the standard deviation of two technical replicates from one experiment.

**Figure S3.** STARD1 KO cells can perform cholesterol import. Pregnenolone measurements from WT and STARD1 KO MA-10 cells treated with or without 22-OH-Chol for 4 hours. Errors bars represent the standard deviation of three technical replicates from one experiment.

**Figure S4.** Cholesterol import with a low dose of rotenone. Relative pregnenolone levels from H295R cells treated with 40 nM Rotenone. Errors bars represent the standard deviation from 4 biological replicates. **p<0.01, unpaired t-test.

**Table S1.** List of antibodies used for western blotting.

## RFERENCES

Arakane, F. et al. (1996) “Steroidogenic acute regulatory protein (StAR) retains activity in the absence of its mitochondrial import sequence: Implications for the mechanism of StAR-action,” Proceedings of the National Academy of Sciences of the United States of America, 93(24), pp. 13731–13736. Available at: 10.1073/pnas.93.24.13731.

Artemenko, I.P. et al. (2001) “Mitochondrial processing of newly synthesized steroidogenic acute regulatory protein (StAR), but not total StAR, mediates cholesterol transfer to cytochrome P450 side chain cleavage enzyme in adrenal cells,” The Journal of Biological Chemistry, 276(49), pp. 46583– 46596. Available at: 10.1074/jbc.M107815200.

Bello-Reuss, E. et al. (2000) “Role of multidrug resistance P-glycoprotein in the secretion of aldosterone by human adrenal NCI-H295 cells,” American Journal of Physiology. Cell Physiology, 278(6), pp. C1256–1265. Available at: 10.1152/ajpcell.2000.278.6.C1256.

Bi, P.Y. et al. (2024) “Cytosolic retention of HtrA2 during mitochondrial protein import stress triggers the DELE1-HRI pathway,” Communications Biology, 7(1), p. 391. Available at: 10.1038/s42003-024-06107-7.

Boos, F. et al. (2019) “Mitochondrial protein-induced stress triggers a global adaptive transcriptional programme,” Nature Cell Biology, 21(4), pp. 442–451. Available at: 10.1038/s41556-019-0294-5.

Bose, H.S., Lingappa, V.R. and Miller, W.L. (2002) “Rapid regulation of steroidogenesis by mitochondrial protein import,” Nature, 417(6884), pp. 87–91. Available at: 10.1038/417087a.

Chen, Y. et al. (2007) “Mitochondrial electron-transport-chain inhibitors of complexes I and II induce autophagic cell death mediated by reactive oxygen species,” Journal of Cell Science, 120(Pt 23), pp. 4155–4166. Available at: 10.1242/jcs.011163.

Choubey, V. et al. (2011) “Mutant A53T α-Synuclein Induces Neuronal Death by Increasing Mitochondrial Autophagy,” The Journal of Biological Chemistry, 286(12), pp. 10814–10824. Available at: 10.1074/jbc.M110.132514.

Corkery-Hayward, M. and Metherell, L.A. (2023) “Adrenal Dysfunction in Mitochondrial Diseases,” International Journal of Molecular Sciences, 24(2), p. 1126. Available at: 10.3390/ijms24021126.

Costa, C.M. et al. (2019) “Levels of cortisol and neurotrophic factor brain-derived in Parkinson’s disease,” Neuroscience Letters, 708, p. 134359. Available at: 10.1016/j.neulet.2019.134359.

Di Maio, R. et al. (2016) “α-Synuclein binds to TOM20 and inhibits mitochondrial protein import in Parkinson’s disease,” Science Translational Medicine, 8(342), p. 342ra78. Available at: 10.1126/scitranslmed.aaf3634.

Djamshidian, A. et al. (2011) “Salivary cortisol levels in Parkinson’s disease and its correlation to risk behaviour,” Journal of Neurology, Neurosurgery, and Psychiatry, 82(10), pp. 1107–1111. Available at: 10.1136/jnnp.2011.245746.

Du, J. et al. (2009) “Dynamic regulation of mitochondrial function by glucocorticoids,” Proceedings of the National Academy of Sciences of the United States of America, 106(9), pp. 3543–3548. Available at: 10.1073/pnas.0812671106.

Epstein, L.F. and Orme-Johnson, N.R. (1991) “Regulation of steroid hormone biosynthesis. Identification of precursors of a phosphoprotein targeted to the mitochondrion in stimulated rat adrenal cortex cells,” The Journal of Biological Chemistry, 266(29), pp. 19739–19745.

Forsström, S. et al. (2019) “Fibroblast Growth Factor 21 Drives Dynamics of Local and Systemic Stress Responses in Mitochondrial Myopathy with mtDNA Deletions,” Cell Metabolism, 30(6), pp. 1040–1054.e7. Available at: 10.1016/j.cmet.2019.08.019.

Franco-Iborra, S. et al. (2018) “Defective mitochondrial protein import contributes to complex I-induced mitochondrial dysfunction and neurodegeneration in Parkinson’s disease,” Cell Death & Disease, 9(11), p. 1122. Available at: 10.1038/s41419-018-1154-0.

Fukumoto, K. et al. (2009) “Detrimental effects of glucocorticoids on neuronal migration during brain development,” Molecular Psychiatry, 14(12), pp. 1119–1131. Available at: 10.1038/mp.2009.60.

Galano, M. and Papadopoulos, V. (2022) “Role of Constitutive STAR in Mitochondrial Structure and Function in MA-10 Leydig Cells,” Endocrinology, 163(8), p. bqac091. Available at: 10.1210/endocr/bqac091.

Gandhi, S. et al. (2012) “Dopamine induced neurodegeneration in a PINK1 model of Parkinson’s disease,” PloS One, 7(5), p. e37564. Available at: 10.1371/journal.pone.0037564.

Granot, Z. et al. (2003) “Proteolysis of normal and mutated steroidogenic acute regulatory proteins in the mitochondria: the fate of unwanted proteins,” Molecular Endocrinology, 17(12), pp. 2461–2476. Available at: 10.1210/me.2003-0074.

Granot, Z. et al. (2007) “Turnover of StAR protein: roles for the proteasome and mitochondrial proteases,” Molecular and Cellular Endocrinology, 265–266, pp. 51–58. Available at: 10.1016/j.mce.2006.12.003.

Hartmann, A. et al. (1997) “Twenty-four hour cortisol release profiles in patients with Alzheimer’s and Parkinson’s disease compared to normal controls: ultradian secretory pulsatility and diurnal variation,” Neurobiology of Aging, 18(3), pp. 285–289. Available at: 10.1016/s0197-4580(97)80309-0.

Jacoupy, M. et al. (2019) “The PINK1 kinase-driven ubiquitin ligase Parkin promotes mitochondrial protein import through the presequence pathway in living cells,” Scientific Reports, 9(1), p. 11829. Available at: 10.1038/s41598-019-47352-9.

Jin, S.M. and Youle, R.J. (2013) “The accumulation of misfolded proteins in the mitochondrial matrix is sensed by PINK1 to induce PARK2/Parkin-mediated mitophagy of polarized mitochondria,” Autophagy, 9(11), pp. 1750–1757. Available at: 10.4161/auto.26122.

Kennedy, B.E., Charman, M. and Karten, B. (2017) “Measurement of Mitochondrial Cholesterol Import Using a Mitochondria-Targeted CYP11A1 Fusion Construct,” Methods in Molecular Biology, 1583, pp. 163–184. Available at: 10.1007/978-1-4939-6875-6_12.

King, S.R. et al. (1999) “Effects of disruption of the mitochondrial electrochemical gradient on steroidogenesis and the Steroidogenic Acute Regulatory (StAR) protein,” The Journal of Steroid Biochemistry and Molecular Biology, 69(1–6), pp. 143–154. Available at: 10.1016/s0960-0760(98)00152-6.

King, S.R. and Stocco, D.M. (1996) “ATP and a mitochondrial electrochemical gradient are required for functional activity of the steroidogenic acute regulatory (StAR) protein in isolated mitochondria,” Endocrine Research, 22(4), pp. 505–514. Available at: 10.1080/07435809609043739.

Lee, J. et al. (2022) “Inhibition of mitochondrial LonP1 protease by allosteric blockade of ATP binding and hydrolysis via CDDO and its derivatives,” The Journal of Biological Chemistry, 298(3), p. 101719. Available at: 10.1016/j.jbc.2022.101719.

Lee-Glover, L.P. and Shutt, T.E. (2024) “Mitochondrial quality control pathways sense mitochondrial protein import,” Trends in endocrinology and metabolism: TEM, 35(4), pp. 308–320. Available at: 10.1016/j.tem.2023.11.004.

Lin, Y.-F. et al. (2016) “Maintenance and propagation of a deleterious mitochondrial genome by the mitochondrial unfolded protein response,” Nature, 533(7603), pp. 416–419. Available at: 10.1038/nature17989.

Lionaki, E. et al. (2022) “Mitochondrial protein import determines lifespan through metabolic reprogramming and de novo serine biosynthesis,” Nature Communications, 13(1), p. 651. Available at: 10.1038/s41467-022-28272-1.

Martinez, T.N. and Greenamyre, J.T. (2012) “Toxin Models of Mitochondrial Dysfunction in Parkinson’s Disease,” Antioxidants & Redox Signaling, 16(9), pp. 920–934. Available at: 10.1089/ars.2011.4033.

Mathis, D. et al. (2024) “Mitochondrial dysfunction results in enhanced adrenal androgen production in H295R cells,” The Journal of Steroid Biochemistry and Molecular Biology, 243, p. 106561. Available at: 10.1016/j.jsbmb.2024.106561.

Mattson, M.P. (2008) “Hormesis defined,” Ageing Research Reviews, 7(1), pp. 1–7. Available at: 10.1016/j.arr.2007.08.007.

Meimaridou, E. et al. (2012) “Mutations in NNT encoding nicotinamide nucleotide transhydrogenase cause familial glucocorticoid deficiency,” Nature Genetics, 44(7), pp. 740–742. Available at: 10.1038/ng.2299.

Naresh, N.U. and Haynes, C.M. (2019) “Signaling and Regulation of the Mitochondrial Unfolded Protein Response,” Cold Spring Harbor Perspectives in Biology, 11(6), p. a033944. Available at: 10.1101/cshperspect.a033944.

Odermatt, A., Strajhar, P. and Engeli, R.T. (2016) “Disruption of steroidogenesis: Cell models for mechanistic investigations and as screening tools,” The Journal of Steroid Biochemistry and Molecular Biology, 158, pp. 9–21. Available at: 10.1016/j.jsbmb.2016.01.009.

Parmar, J., Key, R.E. and Rainey, W.E. (2008) “Development of an adrenocorticotropin-responsive human adrenocortical carcinoma cell line,” The Journal of Clinical Endocrinology and Metabolism, 93(11), pp. 4542–4546. Available at: 10.1210/jc.2008-0903.

Prasad, M. et al. (2012) “Mitochondrial 3β-Hydroxysteroid Dehydrogenase Enzyme Activity Requires Reversible pH-dependent Conformational Change at the Intermembrane Space,” The Journal of Biological Chemistry, 287(12), pp. 9534–9546. Available at: 10.1074/jbc.M111.333278.

Prasad, R. et al. (2014) “Oxidative stress and adrenocortical insufficiency,” The Journal of Endocrinology, 221(3), pp. R63–R73. Available at: 10.1530/JOE-13-0346.

Ravipati, P. et al. (2026) “Protection Against Cellular Toxicity from Rotenone Treatment by the Neuroprotective, Novel Multifunctional Antiparkinsonian Drug D-512,” Journal of Personalized Medicine, 16(2), p. 115. Available at: 10.3390/jpm16020115.

Sierra, A. (2004) “Neurosteroids: the StAR protein in the brain,” Journal of Neuroendocrinology, 16(9), pp. 787–793. Available at: 10.1111/j.1365-2826.2004.01226.x.

Skogar, O. et al. (2011) “Diurnal salivary cortisol concentrations in Parkinson’s disease: increased total secretion and morning cortisol concentrations,” International Journal of General Medicine, 4, pp. 561–569. Available at: 10.2147/IJGM.S20875.

Slominski, R. et al. (2020) “Extra-adrenal glucocorticoid biosynthesis: Implications for autoimmune and inflammatory disorders.,” Genes and immunity, 21(3), pp. 150–168. Available at: 10.1038/s41435-020-0096-6.

Soares, N.M. et al. (2019) “Cortisol levels, motor, cognitive and behavioral symptoms in Parkinson’s disease: a systematic review,” Journal of Neural Transmission, 126(3), pp. 219–232. Available at: 10.1007/s00702-018-1947-4.

Song, J., Herrmann, J.M. and Becker, T. (2021) “Quality control of the mitochondrial proteome,” Nature Reviews Molecular Cell Biology, 22(1), pp. 54–70. Available at: 10.1038/s41580-020-00300-2.

Štefunková, N. et al. (2023) “Comparison of the Effect of BPA and Related Bisphenols on Membrane Integrity, Mitochondrial Activity, and Steroidogenesis of H295R Cells In Vitro,” Life, 14(1), p. 3. Available at: 10.3390/life14010003.

Tack, L.J.W. et al. (2016) “Effects of Glucocorticoids on Bone: What we can Learn from Pediatric Endogenous Cushing’s Syndrome,” Hormone and Metabolic Research = Hormon- Und Stoffwechselforschung = Hormones Et Metabolisme, 48(11), pp. 764–770. Available at: 10.1055/s-0042-117721.

Urban, R.C. and Cotlier, E. (1986) “Corticosteroid-induced cataracts,” Survey of Ophthalmology, 31(2), pp. 102–110. Available at: 10.1016/0039-6257(86)90077-9.

Vasquez, V. et al. (2024) “Mitochondria-targeted oligomeric α-synuclein induces TOM40 degradation and mitochondrial dysfunction in Parkinson’s disease and parkinsonism-dementia of Guam,” Cell Death & Disease, 15(12), p. 914. Available at: 10.1038/s41419-024-07258-5.

Wang, T. and Rainey, W.E. (2012) “Human Adrenocortical Carcinoma Cell Lines,” Molecular and Cellular Endocrinology, 351(1), pp. 58–65. Available at: 10.1016/j.mce.2011.08.041.

Yang, Y. et al. (2019) “PRIMARY ALDOSTERONISM: KCNJ5 MUTATIONS AND ADRENOCORTICAL CELL GROWTH,” Hypertension (Dallas, Tex.2: 1979), 74(4), pp. 809–816. Available at: 10.1161/HYPERTENSIONAHA.119.13476.

Zhang, Q. et al. (2018) “The mitochondrial unfolded protein response is mediated cell-non-autonomously by retromer-dependent Wnt signaling,” Cell, 174(4), pp. 870–883.e17. Available at: 10.1016/j.cell.2018.06.029.

Zhao, T. et al. (2020) “Skeletal Phenotypes Due to Abnormalities in Mitochondrial Protein Homeostasis and Import,” International Journal of Molecular Sciences, 21(21), p. 8327. Available at: 10.3390/ijms21218327.

Zhao, Y. et al. (2012) “Mono-(2-ethylhexyl) phthalate affects the steroidogenesis in rat Leydig cells through provoking ROS perturbation,” Toxicology in vitro: an international journal published in association with BIBRA, 26(6), pp. 950–955. Available at: 10.1016/j.tiv.2012.04.003.

