## Supplemental Table for "Altered Steroidogenesis in Response to Impaired Mitochondrial Protein Homeostasis"

### Table S1.

| Target | Source | Dilution |
|---|---|---|
| **STARD1 (rabbit)** | StAR D10H12<br>Cell Signalling, 8449S | 1:250 – 1:1000 |
| **Beta-actin (mouse)** | Sigma Aldrich, A5316 | 1:3000 |
| **MFN1 (rabbit)** | Cell Signalling, 14739 | 1:1000 |
| **TOM20 (mouse)** | Santa Cruz, SC-17764 | 1:1000 |
| **OPA1 (mouse)** | BDBioscience, 612607 | 1:1000 |
| **SLP-2 (rabbit)** | Alexis Biochem, 210-940-C100 | 1:1000 |
| **GFP (rabbit)** | Abcam, ab290 | 1:1000 |
| **Alpha-synuclein (rabbit)** | Abcam, ab138501 | 1:1000 |
| **FLAG epitope tag (goat)** | Novus Biologicals, NB600-344 | 1:1000 |
| **Anti-mouse HRP** | Invitrogen, 31430 | 1:10 000 |
| **Anti-rabbit HRP** | Cell Signalling, 7074S | 1:10 000 |
| **Anti-goat HRP** | Abcam, ab97110 | 1:10 000 |
