## Supplemental Figures for "Altered Steroidogenesis in Response to Impaired Mitochondrial Protein Homeostasis"

#### Figure S1.

A)

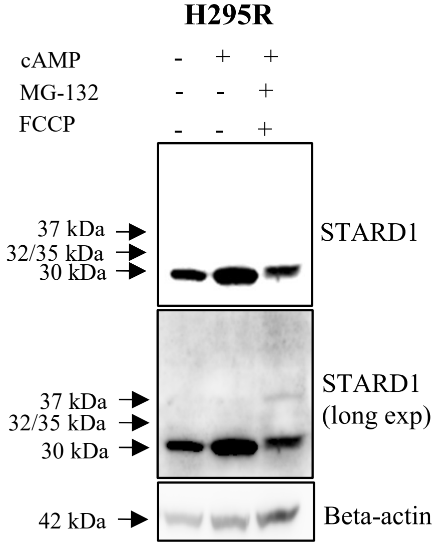


B)

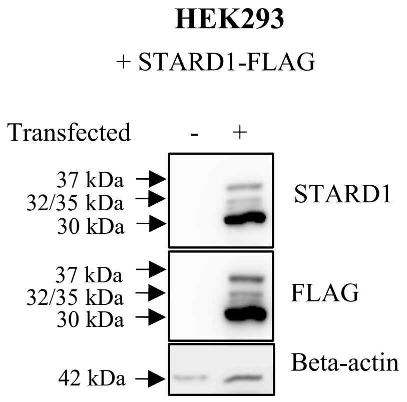

### Figure S2.

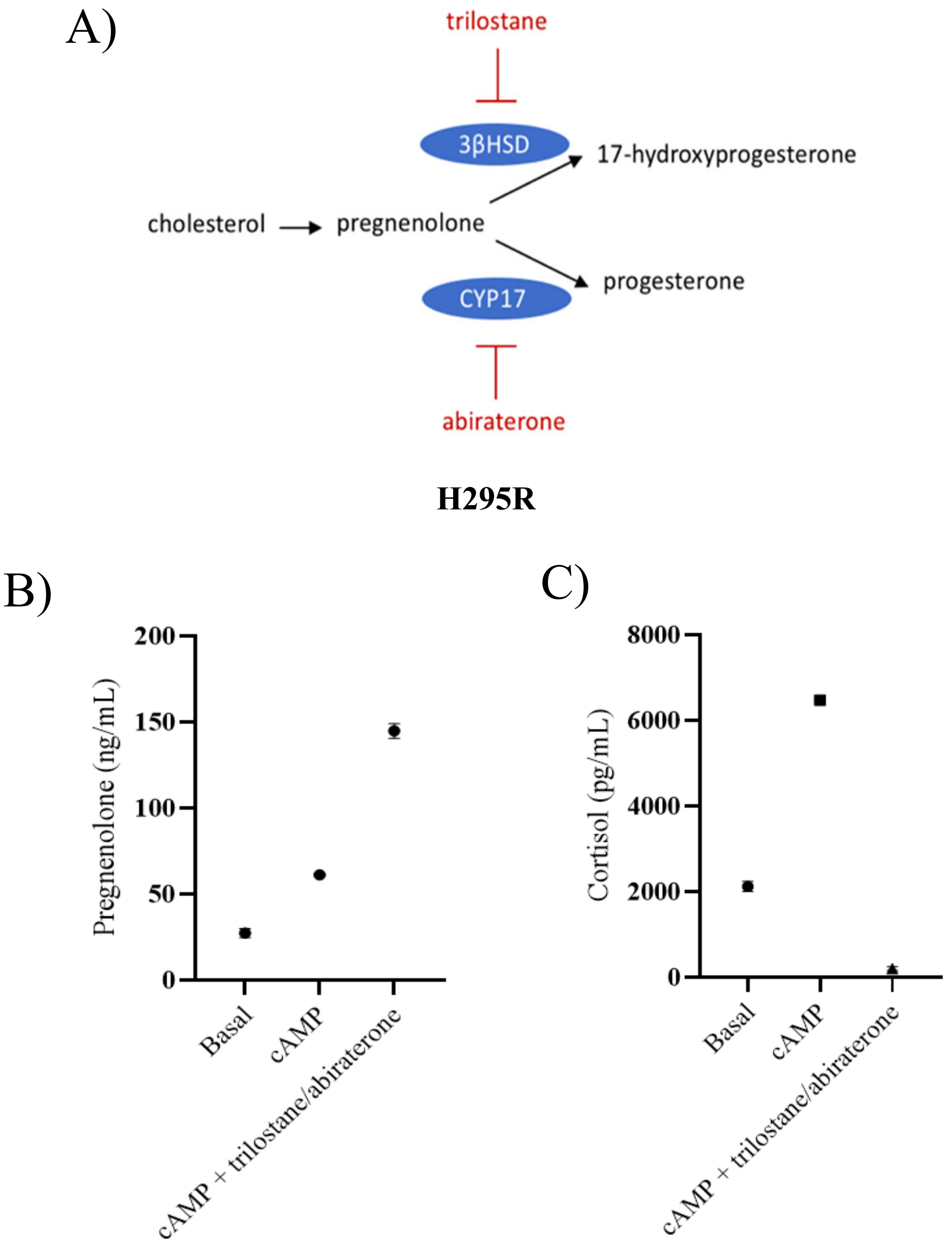
A)
trilostane
3βHSD
17-hydroxyprogesterone
cholesterol
pregnenolone
progesterone
CYP17
abiraterone
H295R
B)
Pregnenolone (ng/mL)
200
150
100
50
0
Basal
cAMP
cAMP + trilostane/abiraterone
C)
Cortisol (pg/mL)
8000
6000
4000
2000
0
Basal
cAMP
cAMP + trilostane/abiraterone

### Figure S3.

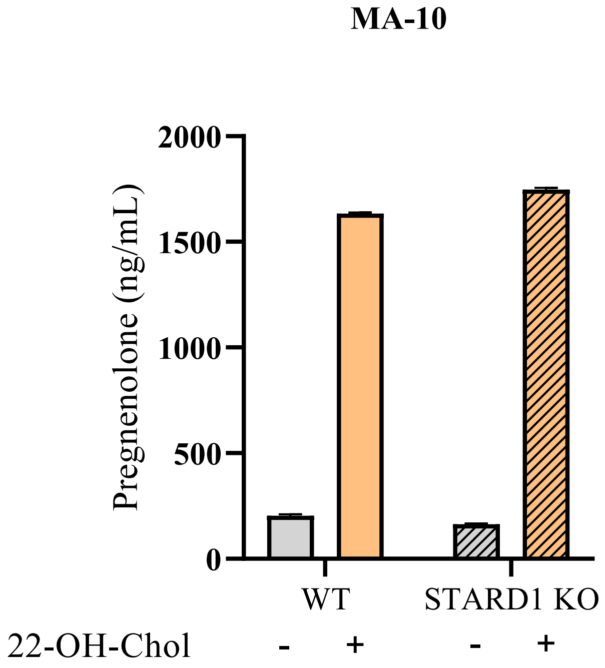
MA-10
2000
1500
1000
500
0
Pregnenolone (ng/mL)
WT
STARD1 KO
22-OH-Chol
-
+
-
+

### Figure S4.

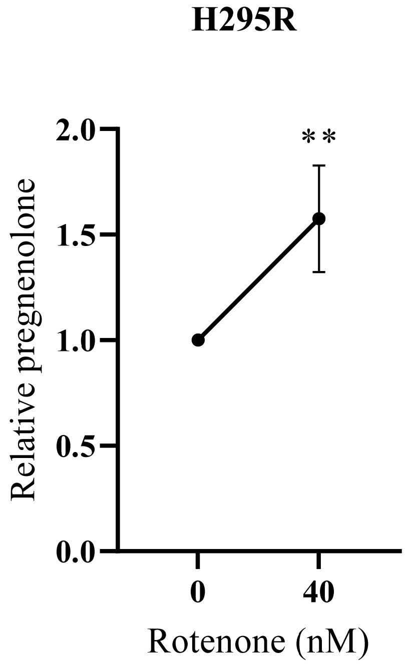
